# Assessing sampling sufficiency of network metrics using bootstrap

**DOI:** 10.1101/080655

**Authors:** Grasiela Casas, Vinicius A.G. Bastazini, Vanderlei J. Debastiani, Valério D. Pillar

## Abstract

Sampling the full diversity of interactions in an ecological community is a highly intensive effort. Recent studies have demonstrated that many network metrics are sensitive to both sampling effort and network size. Here, we develop a statistical framework, based on bootstrap resampling, that aims to assess sampling sufficiency for some of the most widely used metrics in network ecology, namely connectance, nestedness (NODF-nested overlap and decreasing fill) and modularity (using the QuaBiMo algorithm). Our framework can generate confidence intervals for each network metric with increasing sample size (i.e., the number of sampled interaction events, or number of sampled individuals), which can be used to evaluate sampling sufficiency. The sample is considered sufficient when the confidence limits reach stability or lie within an acceptable level of precision for the aims of the study. We illustrate our framework with data from three quantitative networks of plant and frugivorous birds, varying in size from 16 to 115 species, and 17 to 2,745 interactions. These data sets illustrate that, for the same dataset, sampling sufficiency may be reached at different sample sizes depending on the metric of interest. The bootstrap confidence limits reached stability for the two largest networks, but were wide and unstable with increasing sample size for all three metrics estimated based on the smallest network. The bootstrap method is useful to empirical ecologists to indicate the minimum number of interactions necessary to reach sampling sufficiency for a specific network metric. It is also useful to compare sampling techniques of networks in their capacity to reach sampling sufficiency. Our method is general enough to be applied to different types of metrics and networks.

## Introduction

Understanding how species interact with one another is an important and challenging task for ecologists and evolutionary biologists. Over the recent decades the rise and rapid development of network science (Albert and Barabási 2002) has allowed ecologists to improve our understanding of species interactions and their underlying ecological and evolutionary mechanisms. Network studies have grown extraordinarily in the past few decades, and currently they have mainly focused on the analyses of network structure and robustness (Bascompte and Jordano 2007, Miranda et al. 2013).

Some of the most commonly used metrics to describe the structure of ecological networks are connectance, nestedness, modularity, and robustness (Dormann et al. 2009, Miranda et al. 2013). These structural properties have been demonstrated to be associated with community stability and ecosystem functioning (Bascompte et al. 2006, Takimoto and Suzuki 2016). Empirical evidence suggest that most mutualistic networks are nested (Bascompte et al. 2003), and that a highly connected and nested architecture promotes community stability in mutualistic networks. However, in trophic networks this stability is enhanced with compartmented (modular) and weakly connected architectures (Thébault and Fontaine 2010). A modular pattern sometimes prevents the spread of perturbations across the network (Stouffer and Bascompte 2011).

Nevertheless, sampling may influence the detection of network structures. Most network metrics are sensitive to sampling effort and network size (Dormann et al. 2009). Studying pollination networks, Olesen et al. (2007) found a relationship between network size and nestedness and modularity. In general, if network size in seed dispersal increases, connectance decreases (Mello et al. 2011). Bascompte et al. (2003) also found, for plant-frugivore and plant-pollinator networks, that above a size of 50 species, all networks were significantly nested. Consequently, studies having low sampling effort need to be interpreted with caution (Rivera-Hutinel et al. 2012), for low sampling effort can influence the network pattern detected (Vizentin-Bugoni et al. 2015). However, sampling the full diversity of interactions is a highly intensive effort, and ecologists have now come to realize that most networks published to date may be under-sampled. Chacoff et al. (2012) found that, despite a large sampling effort, their pollination network was under-sampled, as they detected less than 60% of the potential interactions. Seed dispersal mutualistic networks may also be insufficiently sampled, since most of the published data sets describe small networks.

Although a robust and well-designed sampling procedure is essential for the quality of data, the optimality of sample size and/or intensity effort depends on the objectives of the study (Orloci and Pillar 1991). A census of the species or interactions in a given area may not be necessary to detect a specific pattern in a network. Similarly, the number of interactions needed to reach sampling sufficiency will be different according to the network metrics and the different types of taxonomic groups within mutualistic and antagonistic networks.

To evaluate the accuracy of the sampling procedure in networks, the approach used is manipulating data after sampling. In this regard, the common analyses applied in the literature for mutualistic networks are rarefaction and accumulation curve analyses (Nielsen and Bascompte 2007, Chacoff et al. 2012, Rivera-Hutinel et al. 2012, Jordano 2015). For instance, Martinez et al. (1999) evaluated the relationships between food-web properties and richness among taxonomic webs and trophic webs using Monte Carlo simulations and confidence intervals. Here, we developed a statistical framework that aims to assess sampling sufficiency for some of the most used metrics in network ecology based on bootstrap resampling.

The bootstrap method (Efron 1979, Efron and Tibshirani 1993) is based on the idea that the distribution of observed values in a sample is the best indicator of their distribution in the sampling universe from which the sample was taken. Our framework is similar to that of Martinez et al. (1999): both, Monte Carlo simulations and bootstrap resampling, are based on repeated sampling. However, while Monte Carlo simulation involves randomization or reshuffling of the data, our resampling is made according to the bootstrap method with replacement, ultimately mimicking the resampling of the sampling universe. Our method is intended to answer the following question: how many interaction events or number of individuals are necessary to be sampled in order to reach stability for a given network metric? We showcase our approach estimating sampling sufficiency for nestedness, modularity, and connectance for three empirical mutualistic networks that widely ranged in size.

## Methods

### Bootstrap resampling technique

We adapted the method of bootstrap resampling from Pillar (1998) to assess sampling sufficiency for network metrics. Our framework is a resampling technique that can generate confidence intervals for each network metric with increasing sample size (i.e., the number of interaction events sampled or the number of observed individuals potentially interacting), which can be used to evaluate sampling sufficiency (Manly 1992, Pillar 1998). The observed values in a sample are taken as “a pseudo sampling universe”, the best available representation of the actual sampling universe from which the sample was taken. Each new sample obtained by resampling with replacement the sample is a “bootstrap sample”. The algorithm is the following:

1. Randomly select a bootstrap sample of *n*_*k*_ interaction events with replacement from the observed sample (pseudo sampling universe) with *n* interaction events;
2. Compute the network metric of interest (*θ*_*k*_) for the bootstrap sample and stores the resulting value;
3. Repeat steps 1 and 2 a large number of times (say 1,000 times);
4. Sort the values of *θ*_*k*_ from the smallest to the largest value. Based on this ordering, delimits the confidence limits for a given specified probability α. For example, with 1,000 bootstrap samples and a probability α of 0.05, the lower confidence limit at a given sample size will be the value of *θ*_*k*_ at the 25^th^ position and the upper limit will be the value of *θ*_*k*_ at the 976^th^ position.
5. Repeat steps 1, 2, 3, and 4 for a new bootstrap sample size *n*_*k*_ + δ, where δ is an increase in sample size, repeating the process up to sample size of *n* interaction events.

Resampling data according to the bootstrap method will create a frequency distribution for the network metric of interest in samples with increasing size, mimicking the resampling of the sampling universe. The sample is considered sufficient within the range of sample sizes evaluated when the confidence limits reach stability or lie within an acceptable level of precision for the objectives of the study (Pillar 1998). Stability of the confidence limits indicates that with samples larger than a certain size there is no further gain in precision for the estimation of the analyzed network metric. We also applied the same algorithm considering the number of captured individuals, instead of the number of interaction, as sampling units. Our aim here was to investigate how many birds were necessary to reach sufficiency for each network metric.

### Network metrics

We assessed sampling sufficiency for three commonly used network metrics:

1. Connectance (C), which is the proportion of realized links in a network relative to the possible number of links (Dunne et al. 2002), with values ranging from 0 to 1. For bipartite networks it is calculated as C=L/(I × J), were L is the number of realized links; I and J are the number of species of each part, e.g., plants and animals. Connectance only distinguishes whether links are present or absent (unweighted, binary links), and the information about interaction frequencies is lost.
2. Nestedness is characterized by a core of highly connected species (generalists) that interact mainly with each other, and a group of specialist species that interact mainly with the generalist species (Bascompte et al. 2003). We used NODF (nested overlap and decreasing fill) algorithm proposed by Almeida-Neto et al. (2008), which corrects biases resulting from matrix fill and matrix dimensions. Similar to connectance, information about interaction frequencies is lost. NODF ranges from 0 (non-nestedness) to 100 (perfect nesting).
3. Modularity is characterized by the degree to which there are groups of nodes (species) that interact more among each other than with other groups (modules) in a network (Guimera and Amaral 2005). We assessed modules using the QuaBiMo algorithm that computes modules in quantitative bipartite networks, based on a hierarchical representation of species link weights and optimal allocation to modules (Dormann and Strauss 2013). It ranges between 0 (random network with no modules) to 1 (maximum modularity).

The method described here has been implemented in R (R Core Team 2013). Network metrics were calculated using the package Bipartite (Dormann et al. 2008). The bootstrap function and a script with an example are available as Supplementary material Appendices 2 and 3, and Table A4.

### Examples from Mutualistic Networks

We illustrate our framework with data from three quantitative networks of plant and frugivorous birds (Table 1). Network size in each dataset varied from 16 to 115 species and 17 to 2,745 interactions events. Despite the fact that the original datasets contained quantitative data, we used two unweighted metrics to assess sampling sufficiency (connectance and NODF) that only distinguish whether links are present or absent (binary links), and one weighted (quantitative) metric (QuaBiMo algorithm). However, it is not possible to use unweighted (binary) data with the bootstrap method because the frequency of interactions is necessary for the resampling procedure.

**Table 1.**
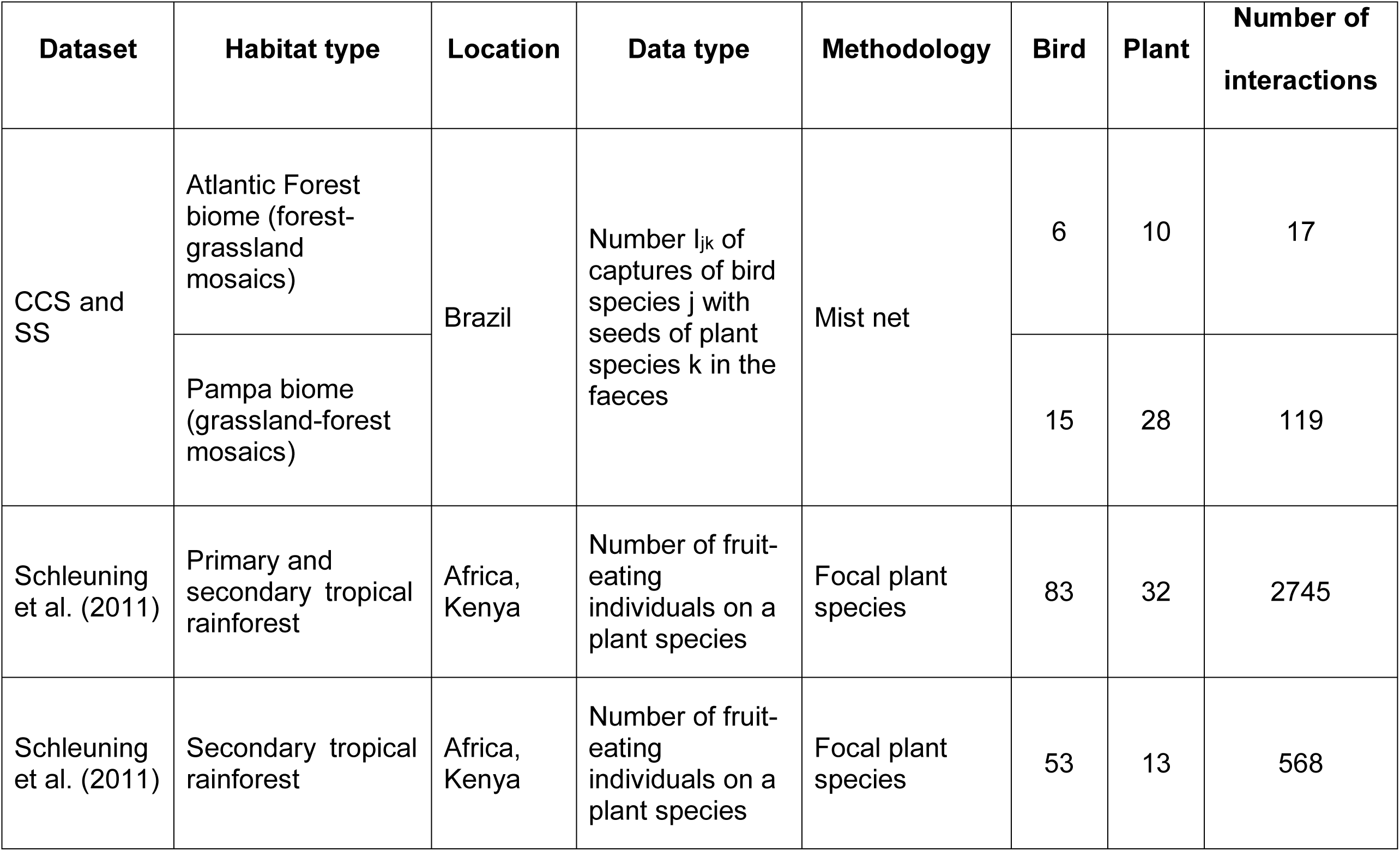
Description of the plant-frugivore networks datasets used with the bootstrap resampling technique.

In two networks, collected by ourselves (named CCS and SS; additional details for methods in Supplementary material Appendix 1), the birds were captured with mist-nets, and then placed into fabric bags to collect their faeces from bags. The seeds found in faecal samples were identified to the species level, when possible, to build an interaction matrix between birds and the plant that they consumed, with the number of interaction events (the number of times a specific bird species was captured with seeds of a specific plant species found in the faeces).

For another test case we used the plant-frugivorous birds network described by Schleuning et al. (2011) (data available from the Interaction Web Database), which was built based on the observation of focal plants, comprising primary and secondary forests and various vegetation strata (Table 1). We used this network as an example of a large quantitative network. To record bird species feeding on each focal plant species, frugivorous bird visits were recorded at each focal plant individual. The interaction frequency was defined as the number of fruit-eating individuals on a plant species independent of fruit handling. We also used, as a separated network, only the data collected in secondary forest areas in Schleuning et al. (2011) (Table 1). The aim was test if the stability for these metrics is reached with a lower sampling effort compared to the entire network of Schleuning et al. (2011), that probably has a high interaction diversity for comprising primary and secondary forests.

To obtain the bootstrap sample (algorithm step 1), for SS, we started with *n*_*k*_ = 10 interaction events with replacement, and we repeated the resampling procedure 1,000 times (algorithm step 3). We then increased sample size by five interaction events (δ), and the process was repeated with *n*_*k*_ + δ up to the maximum number of *n* events. For the smallest network CCS (with only 17 interactions events), we started with *n*_*k*_ = 7 and used δ = 1. For the largest network of Schleuning et al. (2011) (2,745 interactions events) we used *n*_*k*_ = 30 and δ = 50, and for the data collected in secondary forest areas in Schleuning et al. (2011) (with 568 interaction events) we used *n*_*k*_ = 10 and δ = 10. In a second analysis, we used as sample size the actual number of birds captured in our own datasets (CCS and SS), with the same *n*_*k*_ and δ than the interaction events. For all test case, we used 95% confidence intervals based on 1000 resampling interactions at each sample size.

## Results

The bootstrap confidence limits reached stability for the two largest networks for connectance, nestedness, and for modularity within the analyzed sample sizes. Therefore, samples were considered sufficient for these metrics (Fig. 1). The sample of the smallest network (CCS) was not sufficient for any of the analyzed metrics, since the confidence limits were wide and unstable with increasing sample size up to 17 interaction events. For modularity of the CCS network, e.g., the median with the maximum interactions events (17) was expected in 95% of the cases to lie between 0.46 and 0.75, differing by 0.29. This difference is too wide to be indicative of sampling sufficiency compared to the other networks (see detailed results in Supplementary material Appendix Table A1).

**Figure 1.**
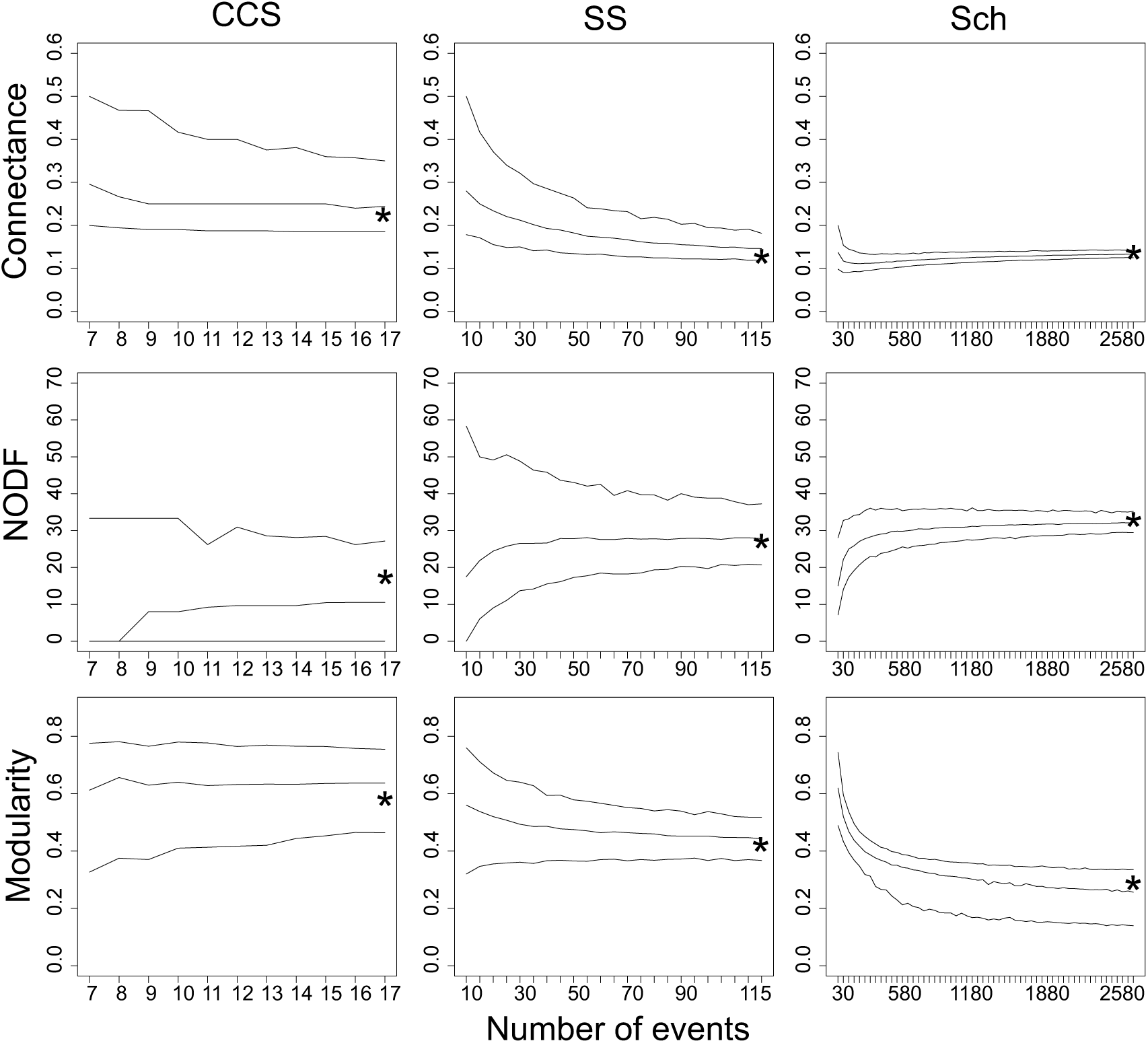
Observed value (star), median and confidence limits of Connectance, Nestedness (NODF) and Modularity metrics obtained by resampling with replacement method using three quantitative mutualistic networks (plants and frugivore birds) and number of interaction events as sample size. The 95% confidence intervals were set based on 1000 resampling interactions at each sample size. See Table 1 for detailed information of networks.

The confidence limits for connectance generated by the bootstrap for the second smallest network (SS) reached stability with sample sizes larger than 90 interaction events, but the confidence limits for nestedness became stable with sample sizes larger than about 60 events (Fig. 1). This means that increasing the sample size beyond 60 interaction events did not add new information that could affect the precision of the bootstrap estimation of nestedness. For the largest network (Schleuning et al. 2001), the stability of confidence limits for connectance, nestedness and modularity was reached, respectively, with sample sizes larger than about 280, 580, and 1000 interaction events. When we considered, as a separated network, only the data collected in secondary forest areas in Schleuning et al. (2011) (with 568 interaction events), we found different results: the bootstrap confidence limits reached stability for the metrics connectance, nestedness and modularity, respectively, with sample sizes larger than about 100, 150 and 230 interaction events (see Supplementary material Appendix Fig. A1).

The results were similar when we considered captured bird individuals as sampling units (Fig. 2). Again, the smallest network (CCS) did not present sufficiency for any of the analyzed metrics, since the confidence limit values were wide and unstable with increasing sample size up to 14 captured individuals. The SS network was considered sufficient for all analyzed metrics, as the bootstrap confidence limits reached stability (Fig. 2; detailed results in Supplementary material Appendix Table A1). Similar results were expected in these cases, for the number of individual consumers were very similar to the number of interaction events because we captured most bird individuals with only one plant species in its faeces. Consequently, the matrix using the number of events and the number of individuals captured in our data were very similar (see Supplementary material Appendix 1 Table A2 for CCS matrix with number of events as sample size, and Table A3 for CCS with number of birds captured as sample size). Despite the similarity in these results, we show and discuss them because the data available for most networks in literature and online databanks unfortunately only allow the extraction of interaction events.

**Figure 2.**
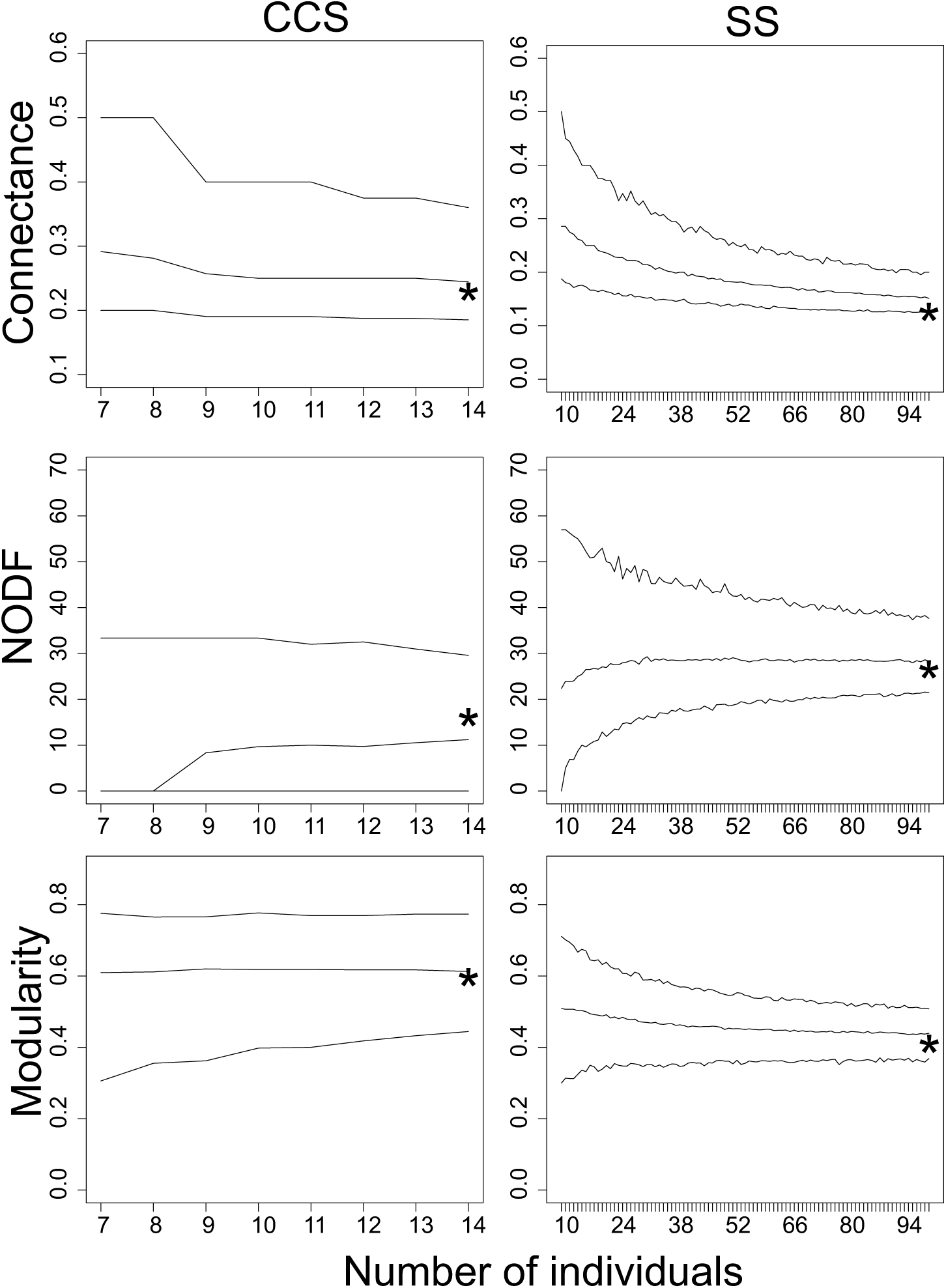
Observed value (star), median and confidence limits of Connectance, Nestedness (NODF) and Modularity metrics obtained by resampling with replacement method using two quantitative mutualistic networks (plants and frugivore birds) and number of bird individuals captured as sample size. The 95% confidence intervals are set based on 1000 resampling interactions at each sample size. See Table 1 for detailed information of seed-dispersal networks (bird and plant) CCS and SS.

## Discussion

We assessed the stability and the precision of the estimated network metrics with the bootstrap method we propose here. Only for the smallest network (with 17 interaction events) the bootstrap confidence limits did not reach stability, but for the others (more than 100 interaction events) the confidence limits reached stability.

The method is general enough to be applied to different types of metrics and networks. However, the type of network metric has to provide a single value at the end of the analysis. For example, modularity involved an optimization method, but we could use it because it gives a modularity *Q* value (Dormann and Strauss 2013). Further, since the aim is to evaluate sampling sufficiency of network metrics, the network must be a quantitative one because data containing the frequency of interactions is necessary for the resampling procedure (interaction events or captured individuals in the test cases). Also, we stress that the bootstrap resampling does not add species and interactions to the network; it only resamples the data with replacement, mimicking the resampling of the presumed sampling universe represented by the observed sample.

With the bootstrap method we are looking for the effect of sampling bias on network metrics. In a different way, previous studies compared different fieldwork sampling techniques and investigated to which extent their conclusions (structural properties of network) were influenced by the way samples were collected. For example, Gibson et al. (2011) analyzed the potential bias in network metrics when using time-based observations or transects in plant-pollinator networks, with rarefaction analysis and null models approach. Analogously, the bootstrap method can be used to compare two methodologies in terms of sampling sufficiency. In seed dispersal networks between plants and birds, e.g., the sampling hours or number of observed plant individuals as sample size (through transect or focal-plant methodologies) can be compared with the number of bird individuals captured (with mist net) that need to be sampled in order to reach stability for each network metric.

We assessed sampling sufficiency with the bootstrap method using interaction events as sampling units with three mutualistic networks that differed regarding sampling techniques. In addition, we used the captured bird individuals as sampling units to assess sufficiency for the networks we have collected. The potential advantage of using individual data over interaction events is that the first is more independent in comparison to the latter. Independence between sampling units is often an important assumption in data analysis, and the accuracy of the bootstrap method may be affected by lack of independence (Efron and Tibshirani 1993). However, in spite of the advantage of using individuals as sampling units, the data available for most networks in literature and online databanks unfortunately only allow the extraction of interactions events.

In our study, we observed a variation of the median value of the metric generated by bootstrap resampling, mainly in small samples. We expected that the median would remain stable throughout the process, and only the confidence interval would change with increasing sample size. Since most network metrics are sensitive to sampling effort and network size (Dormann et al. 2009), probably these metrics are biased, causing this variation in the median value of the metric. Further research, using simulated networks with known properties and dimensions, is needed to evaluate how accurate these network metrics are.

Our results suggest an important point: sampling sufficiency can be reached at different sample sizes for the same dataset depending on the metric of interest. Nielsen and Bascompte (2007), analyzing the sensitivity of connectance and nestedness metrics to variation in sampling effort, also suggested that sampling intensity does not affect all network metrics in the same way, and that nestedness tends to stabilize rapidly with increasing sampling effort. Some confidence limits generated by the bootstrap reached stability with less than 100 interaction events, meaning that sampling more interaction events probably would have not significantly affected the estimation of this network pattern.

However, it has been pointed out that studies of interactions should come from a robust and well-designed sampling procedure, mainly due to the influence of limited sampling effort in network properties (Dormann et al. 2009, Vázquez et al. 2009, Chacoff et al. 2012, Vizentin-Bugoni et al. 2015). In our results, even though the bootstrap confidence limits for some network metrics reached stability in networks with less than 50 species, the range of confidence limits for the largest network (Schleuning et al. 2011), with 115 species, was much smaller compared to the other two networks and, consequently, it is considered a more precise sample. Because the study of Schleuning et al. (2011) comprised primary and secondary forests and various vegetation strata, the interactions of this network are heterogeneous (high interaction diversity) and, consequently, the stability for these metrics was reached only with a larger sampling effort compared to the other networks.

The bootstrap method we propose here can help network ecologists by indicating the minimum number of interaction events (or other defined sampling units) necessary to reach sampling sufficiency for a specific network metric. It also allows comparing sampling protocols in terms of effort to reach sampling sufficiency. The concerns on the effect of sampling effort on network metrics in mutualistic (Nielsen and Bascompte 2007, Dorado et al. 2011, Chacoff et al. 2012, Vizentin-Bugoni et al. 2015) and food webs (Goldwasser and Roughgarden 1997, Martinez et al. 1999, Banašek-Richter et al. 2004) have grown in the last few years. We believe that our method is a significant contribution to assess sampling sufficiency in network ecology.

## Acknowledgements

We thank Programa de Pós-Graduação em Ecologia – Universidade Federal do Rio Grande do Sul and the Conselho Nacional de Desenvolvimento Científico e Tecnológico (CNPq) for funding received. This work is part of the project Biodiversity of grasslands and grassland-forest ecotones in Southern Brazil: ecological basis for conservation and sustainable use (SISBIOTA-Federal funding: MCT/CNPq/MEC/CAPES/FNDCT, grant 563271/2010-8). GC thanks CAPES for a scholarship and for the sandwich internship (Ministry of Education of Brazil, Brasília – DF 70.40-020, Brazil), conducted on Pacific Ecoinformatics and Computational Ecology Lab, coordinated by Dr. Neo Martinez, where the first discussion of sampling sufficiency started. VDP has been supported by a CNPq fellowship (grant 307689/2014-0).

## Supplementary material

Table A1 – A4. Fig. A1. Appendix 1 – 3.

